# Ebselen, disulfiram, carmofur, PX-12, tideglusib, and shikonin are non-specific promiscuous SARS-CoV-2 main protease inhibitors

**DOI:** 10.1101/2020.09.15.299164

**Authors:** Chunlong Ma, Yanmei Hu, Julia Alma Townsend, Panagiotis I. Lagarias, Michael Thomas Marty, Antonios Kolocouris, Jun Wang

## Abstract

There is an urgent need for vaccines and antiviral drugs to combat the COVID-19 pandemic. Encouraging progress has been made in developing antivirals targeting SARS-CoV-2, the etiological agent of COVID-19. Among the drug targets being investigated, the viral main protease (M^pro^) is one of the most extensively studied drug targets. M^pro^ is a cysteine protease that hydrolyzes the viral polyprotein at more than 11 sites and it is highly conserved among coronaviruses. In addition, M^pro^ has a unique substrate preference for glutamine in the P1 position. Taken together, it appears that M^pro^ inhibitors can achieve both broad-spectrum antiviral activity and a high selectivity index. Structurally diverse compounds have been reported as M^pro^ inhibitors, with several of which also showed antiviral activity in cell culture. In this study, we investigated the mechanism of action of six previously reported M^pro^ inhibitors, ebselen, disulfiram, tideglusib, carmofur, shikonin, and PX-12 using a consortium of techniques including FRET-based enzymatic assay, thermal shift assay, native mass spectrometry, cellular antiviral assays, and molecular dynamics simulations. Collectively, the results showed that the inhibition of M^pro^ by these six compounds is non-specific and the inhibition is abolished or greatly reduced with the addition of reducing reagent DTT. In the absence of DTT, these six compounds not only inhibit M^pro^, but also a panel of viral cysteine proteases including SARS-CoV-2 papain-like protease, the 2A^pro^ and 3C^pro^ from enterovirus A71 (EV-A71) and EV-D68. However, none of the compounds inhibits the viral replication of EV-A71 or EV-D68, suggesting that the enzymatic inhibition potency IC_50_ values obtained in the absence of DTT cannot be used to faithfully predict their cellular antiviral activity. Overall, we provide compelling evidence suggesting that ebselen, disulfiram, tideglusib, carmofur, shikonin, and PX-12 are non-specific SARS-CoV-2 M^pro^ inhibitors, and urge the scientific community to be stringent with hit validation.

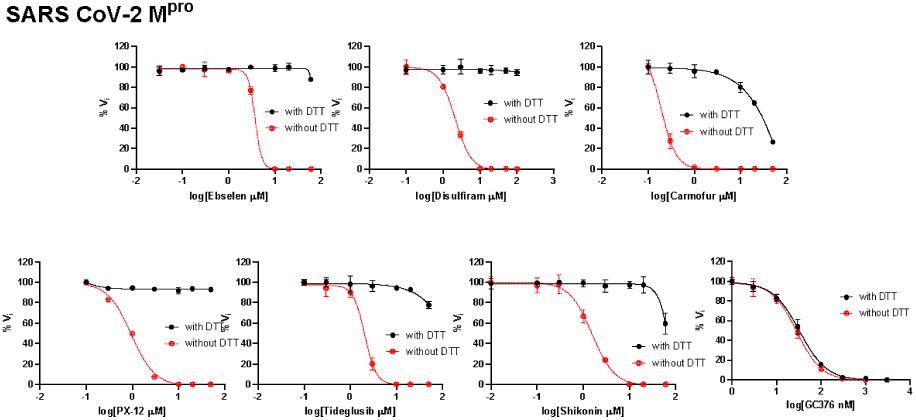

## Introduction

A new coronavirus, SARS-CoV-2, started to circulate among humans in late 2019 and quickly evolved to be a global pandemic. As of August 31, 2020, there have been more than 6 million positive cases with over 183,000 deaths in the United States alone. The origin of SARS-CoV-2 is still under investigation, and the closest strain is the bat coronavirus RaTG13, which shares 96% sequence similarity with SARS-CoV-2.^1^ It is unknown whether SARS-CoV-2 transmitted directly from bats to humans or through an intermediate host.^2^ There is currently no vaccines or antiviral drugs available for SARS-CoV-2. Encouraging progress has been made in vaccine development, and several RNA, DNA, and adenovirus-based vaccine candidates are now in phase III clinical trials.^3^ For small molecule antivirals, remdesivir was granted emergency use authorization in the United States.

SARS-CoV-2 is an enveloped, positive-sense, single-stranded RNA virus that belongs to the betacoronavirus genera, which also includes SARS-CoV, MERS-CoV, HCoV-OC43, and HCoV-HKU1. SARS-CoV-2 shares ∼80% sequence similarity with SARS-CoV. As such, many of the reported antivirals against SARS-CoV-2 were originally developed for SARS-CoV or other related coronaviruses.^4^ SARS-CoV-2 infects ACE2 expressing cells and enters cell through either direct cell surface fusion or endosomal pathway.^5^ For direct cell surface fusion, the host membrane protease TMPRSS2 cleaves the viral spike protein, trigging the viral membrane fusion with the host cell membrane.^6^ For the endosomal entry, cathepsin L mediates the cleavage of viral spike protein.^7^ Once the viral RNA is released in the cytoplasm, it undergoes translation into viral polyproteins pp1a and pp1ab, which are subsequently cleaved by two viral proteases, the main protease (M^pro^), also called 3-chymotrypsin-like protease (3CL^pro^), and the papain-like protease (PL^pro^). The released viral proteins can then assemble to form the viral polymerase RdRp complex to catalyze the replication of viral RNA. Finally, the progeny virions are released from the infected cells through exocytosis and are ready for the next round of infection.

Compounds that interfere with any step in the viral life cycle are expected to inhibit viral replication. Among the list of drug targets pursued for SARS-CoV-2 small molecule antivirals, the viral polymerase RdRp and the protease M^pro^ are the most extensively studied ones. The RdRp inhibitor, remdesivir, received emergency use authorization in the United States. It showed broad-spectrum antiviral activity against multiple coronavirus including SARS-CoV-2, SARS-CoV, and MERS-CoV in cell culture, and it also had in vivo efficacy in SARS-CoV infection mouse model.^8^ EIDD-2801, an RdRp inhibitor, is another promising drug candidate with broad-spectrum antiviral activity.^9^ The M^pro^ is a cysteine protease that cleaves the viral polyprotein at more than 11 sites. It has a unique substrate preference of glutamine at the P1 position of the substrate, while no host protease is known to have such a preference.^10^ As such, the most potent M^pro^ inhibitors such as GC376 and N3 all contain a 2-pyrrolidone substitution in the P1 position as a mimetic of the glutamine in the substrate. Several crystal structures of M^pro^ in complex with inhibitors have been solved, showing that the pyrrolidone forms multiple hydrogen bonds with the His163 and Glu166 side chains and the main chain of Phe140.^11-14^ In addition to the classic pyrrolidone-containing M^pro^ inhibitors, several non-canonical M^pro^ inhibitors have also been reported with both enzymatic inhibition and cellular antiviral activity.^11-12^ In this study, we aim to validate six previously reported M^pro^ inhibitors, ebselen, disulfiram, tideglusib, carmofur, shikonin, and PX-12 (Fig. 1).^12^ Among the six compounds, ebselen is a clinical candidate with anti-inflammatory and anti-oxidant activities. In preclinical studies, ebselen was reported to react with cysteine residues from completely unrelated proteins including the C-terminal domain of the HIV-1 capsid,^15^ the mycobacterium tuberculosis transpeptidase LdtMt2,^16^ glutamate dehydrogenase,^17^ clostridium difficile toxins TcdA and TcdB,^18^ the mycobacterium tuberculosis (Mtb) antigen 85C enzyme,^19^ the hepatitis C virus NS3 helicase,^20^ and many others. Ebselen was also reported to induce protein unfolding for the insulin-degrading enzyme.^21^ Ebselen analogues were synthesized and were found to inhibit both SARS-CoV-2 M^pro^ and PL^pro^.^22^ Disulfiram inhibits a panel of diverse enzymes including methyltransferase,^23^ urease,^24^ and kinase^25^ through reacting with the cysteine residues. A study also showed that disulfiram inhibits the PL^pro^ from SARS-CoV and MERS-CoV with IC_50_ values of 14.2 and 22.7 µM, respectively.^26^ However, the inhibition was completely lost in the presence of 5 mM of β-mercaptoethanol (IC_50_ > 300 µM).^26^ Carmofur inhibits human acid ceramidase by covalently modify the catalytic C143 residue.^27^ PX-12 inhibits tubulin polymerization through cysteine oxidation.^28^ Tideglusib is an irreversible inhibitor of glycogen synthase kinase-3β (GSK-3β).^29^

**Figure 1.**
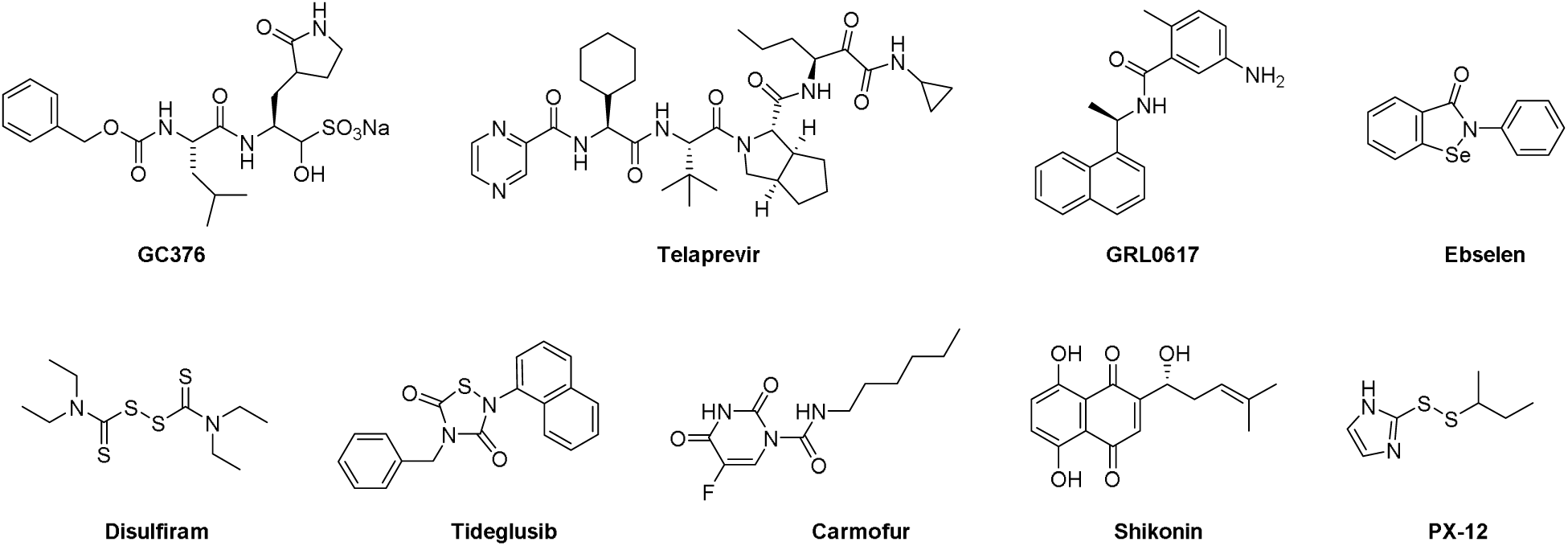
Chemical structures of protease inhibitors investigated in this study.

Ebselen, disulfiram, carmoufur, PX-12, tideglusib, and shikonin were recently reported as SARS-CoV-2 M^pro^ inhibitors with IC_50_ values ranging from 0.67 to 21.39 µM in the FRET-based enzymatic assay.^12^ Among the six compounds, ebselen inhibited SARS-CoV-2 replication with an EC_50_ value of 4.67 ± 0.80 µM in plaque reduction assay. Disulfiram was able to reduce the viral replication by ∼30% in the viral titer reduction assay at 10 µM, while other compounds tideglusib, carmofur, and PX-12 had no significant antiviral effect. Intriguingly, in a follow up study, carmofur was shown to inhibit SARS-CoV-2 viral replication with an EC_50_ of 24.30 µM, and the X-ray crystal structure of M^pro^ with carmofur was solved (PDB: 7BUY).^30^ The description of the enzymatic assay did not specify whether reducing reagent dithiothreitol (DTT) was added or not. It is a standard practice to add DTT or other reducing reagent such as glutathione (GSH) or β-mercaptoethanol (β-ME) in the enzymatic assay of cysteine protease to ensure that the enzyme is in the active form by reducing the catalytic cysteine residue as well as preventing non-specific thiol reactive compounds to covalently modify the catalytic cysteine.

The questions we are trying to address in this study are whether the inhibition of M^pro^ by these compounds is specific and whether their enzymatic inhibition potency IC_50_ values can be used to faithfully predict the cellular antiviral activity. In other words, do the reported IC_50_ values of ebselen, disulfiram, tideglusib, carmofur, shikonin, and PX-12 against SARS-CoV-2 M^pro^ reflect specific enzymatic inhibition, or is it due to non-specific inactivation of the enzyme? For this, we tested these compounds against a panel of related and unrelated viral cysteine proteases, the SARS-CoV-2 PL^pro^, and the 2A protease (2A^pro^) and 3C protease (3C^pro^) from EV-A71 and EV-D68 in a consortium of assays either with or without DTT. Collectively, our results showed that in the absence of DTT, ebselen, disulfiram, tideglusib, carmofur, shikonin, and PX-12 non-specifically inhibit all six viral cysteine proteases including SARS-CoV-2 M^pro^. However, despite their potent inhibition of enzymatic activity of 2A^pro^ and 3C^pro^ from EV-A71 and EV-D68 in the FRET assay in the absence of DTT, none of the compounds showed cellular antiviral activity against EV-A71 and EV-D68. Therefore, it can be concluded that the enzymatic inhibition potency of cysteine protease inhibitors obtained in the absence of DTT cannot be used to predict the cellular antiviral activity. Overall, although there is an immediate need for SARS-CoV-2 antivirals, the scientific community needs to be cautious about the non-specific effect of promiscuous compounds, and secondary assays should be performed at the early stage to triage hits that lack translational potential.

## Results

### The inhibition of SARS-CoV-2 M^pro^, PL^pro^, and EV-A71 and EV-D68 2A^pro^ and 3C^pro^ by ebselen, disulfiram, carmofur, PX-12, tideglusib, and shikonin is DTT-dependent

To dissect the effect of DTT on the enzymatic inhibition of SARS-CoV-2 M^pro^ by ebselen, disulfiram, carmoufur, PX-12, tideglusib, and shikonin, we performed dose response titration in the FRET-based enzymatic assay with and without 4 mM DTT. A known M^pro^ inhibitor, GC376, was included as a positive control. It was found that all compounds inhibited M^pro^ in the absence of DTT (Fig. 2A, red curves), and the IC_50_ values are generally in agreement with previous published results,^12^ except shikonin and PX-12, which showed more than 10-fold difference. However, none of the compounds showed potent inhibition against M^pro^ in the presence of 4 mM DTT (IC_50_ > 25 µM) (Fig. 2A, black curves). Carmofur showed weak inhibition with an IC_50_ value of 28.2 ± 9.5 µM in the presence of DTT (Table 1). In contrast, GC376 showed consistent inhibition against M^pro^ both in the absence and presence of DTT with IC_50_ values of 0.03 µM and 0.03 µM, respectively (Fig. 2A, last column; Table 1). These results suggest that the claimed inhibition of M^pro^ by these six compounds might not be target specific. To test this hypothesis, we next tested these six compounds against five other viral cysteine proteases, among which SARS-CoV-2 PL^pro^, EV-A71 2A^pro^, and EV-D68 2A^pro^ have no sequence similarity as M^pro^, while EV-A71 3C^pro^ and EV-D68 3C^pro^ share similar chymotrypsin-like folding with M^pro^, despite only showing 16.3% and 19.6% sequence similarities as M^pro^. GRL0617 was included as a positive control for SARS-CoV-2 PL^pro^, and telaprevir was included as a positive control for EV-A71 2A^pro^ and EV-D68 2A^pro^. GC376 was used as a positive control for both EV-A71 3C^pro^ and EV-D68 3C^pro^. If the inhibition of M^pro^ by ebselen, disulfiram, carmoufur, PX-12, tideglusib, and shikonin is specific, either in the absence or the presence of DTT, these six compounds are expected to show little or no inhibition against the unrelated SARS-CoV-2 PL^pro^, EV-A71 2A^pro^, and EV-D68 2A^pro^. For the SARS-CoV-2 PL^pro^, we observed similar results as SARS-CoV-2 M^pro^: all compounds displayed potent inhibitory effect in the absence of DTT, while little or no inhibition was observed in the presence of DTT (Fig. 2B). Shikonin was less potent against PL^pro^ than M^pro^ in the absence of DTT with IC_50_ values of 1.5 µM and 55.3 µM, respectively (Table 1). The potency of shikonin increased about two-fold and the IC_50_ values were 55.3 µM and 28.2 µM with and without DTT, respectively (Table 1). As expected, the non-covalent SARS-CoV-2 PL^pro^ inhibitor GRL0617^31^ inhibits SARS-CoV-2 PL^pro^ similarly in the present or the absence of 4 mM DTT (Fig. 2B, last column). For EV-A71 and EV-D68 2A^pro^ and 3C^pro^, we observed similar trends as M^pro^: all six compounds showed potent enzymatic inhibition in the absence of DTT and the inhibition was abolished with the addition of DTT (Figs. 2C-F). In contrast, there is no significant shift of the potency with and without DTT for GC376 in inhibiting EV-A71 3C^pro^ (Fig. 2D) and EV-D68 3C^pro^ (Fig. 2F), and telaprevir in inhibiting EV-A71 2A^pro^ (Fig. 2C) and EV-D68 2A^pro^ (Fig. 2E).

**Table 1.** Enzymatic assay results of protease inhibitors investigated in this study against SARS-CoV-2, EV-A71 and EV-D68 proteases.

|  | SARS-CoV-2 M<sup>pro</sup><br>IC<sub>50</sub> (μM)<br>no DTT/with DTT | SARS-CoV-2 PL<sup>pro</sup><br>IC<sub>50</sub> (μM)<br>no DTT/with DTT | EV-A71 2A<sup>pro</sup><br>IC<sub>50</sub> (μM)<br>no DTT/with<br>DTT | EV-A71 3C<sup>pro</sup><br>IC<sub>50</sub> (μM)<br>no DTT/with DTT | EV-D68 2A<sup>pro</sup><br>IC<sub>50</sub> (μM)<br>no DTT/with<br>DTT | EV-D68 3C<sup>pro</sup><br>IC<sub>50</sub> (μM)<br>no DTT/with DTT |
| --- | --- | --- | --- | --- | --- | --- |
| GC376 | 0.03 ± 0.01/0.03 ± 0.01 | N.T. | N.T. | 0.06 ± 0.02/0.08 ± 0.02 | N.T. | 0.06 ± 0.01/0.05 ± 0.02 |
| Telaprevir | N.T. | N.T. | 1.8 ± 0.9/1.3 ± 0.6 | N.T. | 0.1 ± 0.0/0.2 ± 0.0 | N.T. |
| GRL0617 | N.T. | 1.8 ± 0.2/1.9 ± 0.2 | N.T. | N.T. | N.T. | N.T. |
| Ebselen | 3.7 ± 2.4/>60<br>**0.67 ± 0.09<sup>a</sup>**<br>(reported) | 10.3 ± 8.9/>60 | 5.9 ± 1.1/>60 | 1.2 ± 0.7/>60 | 3.6 ± 1.0/>60 | 0.1 ± 0.0/>60 |
| Disulfiram | 2.1 ± 0.3/>60<br>**9.35 ± 0.18**<br>(reported) | 6.9 ± 4.2/>60 | 11.8 ± 2.1/>60 | 1.0 ± 0.6/>60 | 3.5 ± 0.5/>60 | 0.6 ± 0.1/>60 |
| Tideglusib | 2.1 ± 0.3/>60<br>**1.55 ± 0.30**<br>(reported) | 7.1 ± 1.4/30.4 ± 17.1 | 3.4 ± 2.9/>60 | 1.2 ± 0.1/>60 | 1.3 ± 0.7/>60 | 0.6 ± 0.3/>60 |
| Carmofur | 0.2 ± 0.1/28.2 ± 9.5<br>**1.82 ± 0.06**<br>(reported) | 0.7 ± 0.1/>60 | 12.9 ± 4.5/>60 | 0.4 ± 0.2/>60 | 6.4 ± 1.3/>60 | 0.3 ± 0.0/>60 |
| Shikonin | 1.5 ± 0.3/>60<br>**15.75 ± 8.22**<br>(reported) | 55.3 ± 17.7/28.2 ± 12.5 | 36.0 ± 20.5/>60 | 0.5 ± 0.2/>60 | 37.0 ± 14.2/>60 | 1.2 ± 0.7/>60 |
| PX-12 | 0.9 ± 0.2/>60<br>**21.39 ± 7.06**<br>(reported) | 18.7 ± 2.6/>60 | 16.9 ± 9.2/>60 | 4.1 ± 1.9/>60 | 9.3 ± 4.2/>60 | 1.2 ± 0.2/>60 |

<sup>a</sup>The values shown in bold were reported in reference<sup>12</sup>

**Figure 2.**
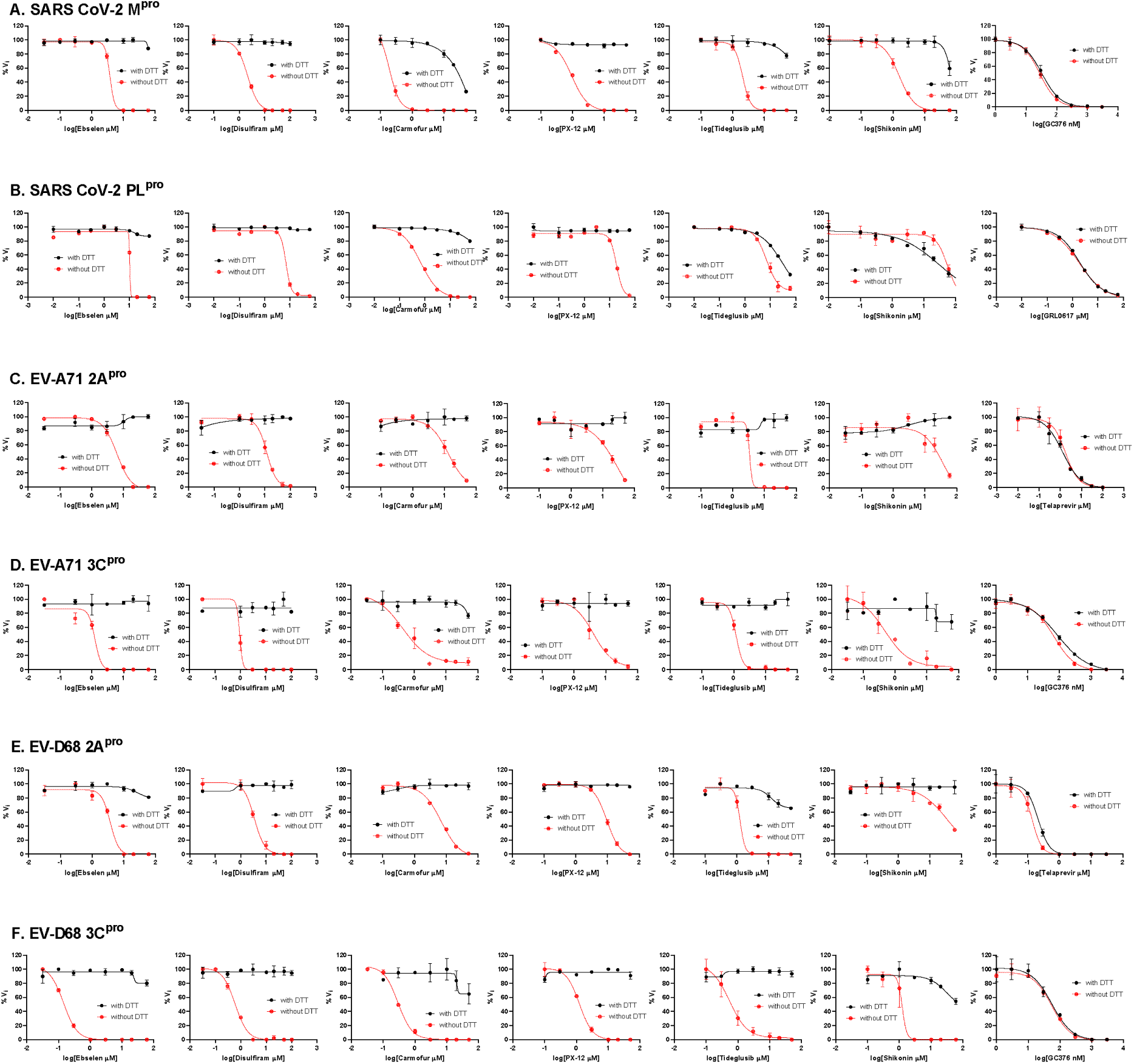
Enzymatic assay of SARS-CoV-2 M^pro^ and PL^pro^, EV-A71 and EV-D68 2A^pro^ and 3C^pro^ against inhibitors investigated in this study. (**A**) SARS-CoV-2 M^pro^; (**B**) SARS-CoV-2 PL^Pro^; (**C**) EV-A71 2A^pro^; (**D**) EV-A71 3C^pro^; (**E**) EV-D68 2A^pro^; and (**F**) EV-D68 3C^pro^. Protease was pre-incubated in their corresponding reaction buffer as described in the method section with various concentrations of protease inhibitors in the presence of 4 mM DTT or in the absence of DTT at 30 °C for 30 min. The enzymatic reaction was initiated by adding the corresponding FRET substrate. The efficacy of these protease inhibitors in the presence of 4 mM DTT or in the absence of DTT was evaluated with a four parameters dose response curve function in prism 8 as described in the method section.

Next, we tested whether another reducing agent glutathione (GSH) could also abolish the inhibitory effect of these promiscuous inhibitors against SARS-CoV-2 M^pro^. For these, ebselen and disulfiram were chosen as representative examples. In the absence of 4 mM DTT or 1 mM GSH, ebselen and disulfiram completely inhibits M^pro^ enzymatic activity at 20 µM (Fig. 3); however, no inhibition was observed for ebselen and disulfiram when either 4 mM DTT or 1 mM GSH was present in the reaction buffer (Fig. 3). In contrast, the inhibition by the positive control GC376 was not affected by the reducing agents DTT or GSH.

**Figure 3.**
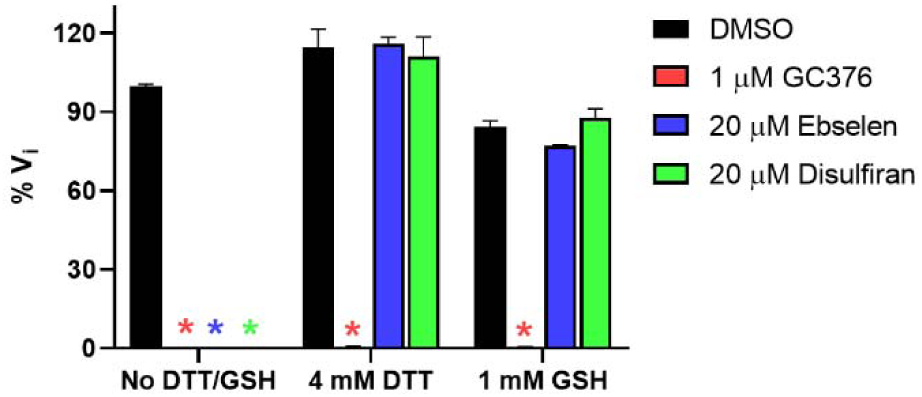
Effect of glutathione (GSH) on the inhibition of ebselen and disulfiram against SARS-CoV-2 M^pro^. 100 nM SARS-CoV-2 M^pro^ protein was pre-incubated in SARS-CoV-2 M^pro^ reaction buffer with the testing protease inhibitors in the absence of DTT or GSH, or in the presence of 4 mM DTT or 1 mM GSH at 30 °C for 30 min. The enzymatic reaction was initiated by adding 10 µM SARS-CoV-2 M^pro^ FRET substrate. The initial enzymatic reaction velocity was measured and normalized to the condition that no protease inhibitor (DMSO) and no DTT/GSH was present in the reaction buffer.

Collectively, the enzymatic assay results suggest that ebselen, disulfiram, carmoufur, PX-12, tideglusib, and shikonin are promiscuous cysteine protease inhibitors that not only inhibit M^pro^ but also five other related and unrelated viral cysteine proteases including SARS-CoV-2 PL^pro^, EV-A71 and EV-D68 2A^pro^ and 3C^pro^ in the absence of DTT, and the inhibition is abolished with the reducing reagent of either DTT or GSH.

### Ebselen, disulfiram, carmofur, PX-12, tideglusib, and shikonin did not bind to SARS-CoV-2 M^pro^ in the presence of DTT in the thermal shift assay

To investigate whether ebselen, disulfiram, carmoufur, PX-12, tideglusib, and shikonin bind directly to M^pro^ or other related and unrelated cysteine proteases, we performed thermal shift binding assay. In the thermal shift binding assay, a temperature gradient is applied to denature a protein in the presence of a fluorescence dye. When protein unfolds, the hydrophobic region is exposed to the fluorescence dye and increased fluoresce signal is observed. Specific binding of a small-molecule to the native state of a protein usually stabilizes the protein, leading to a shift of the melting temperature (ΔT_m_).^32-34^ Here we measured T_m_ change upon addition of these six compounds in the presence or absence of 4 mM DTT against six viral cysteine proteases including M^pro^. All compounds were tested at 40 µM except shikonin, which was tested at 10 µM as it quenches the SYPRO orange dye fluorescence signal at 40 µM. Compounds were pre-incubated with 3 µM protease in its corresponding enzymatic reaction buffer at 30 °C for 30 min, then a 20 to 90 °C temperature gradient was applied and T_m_ was calculated. As expected, for the positive controls, significant T_m_ increase was observed in the binding of GC376 to SARS-CoV-2 M^pro^ (Fig. 4A), EV-A71 3C^pro^ (Fig. 4D) and EV-D68-3C^pro^ (Fig. 4F). Binding of GRL0617 to SARS-CoV-2 PL^Pro^ also led to significant stabilization (Fig. 4B). Similarly, binding of telaprevir to EV-A71 2A^pro^ and EV-D68 2A^pro^ increased the T_m_ (Figs. 4C and 4E). Importantly, there was no different in the T_m_ shift with and without DTT for these positive controls (Figs. 4A-F). In contrast, in the presence of 4 mM DTT, no T_m_ change was observed with the addition of ebselen, disulfiram, carmofur, PX-12, tideglusib, and shikonin to all six proteases, suggesting none of the compounds bind to any of these proteases (Figs. 4A-F). In the absence of DTT, upon the addition of these six compounds, a decrease of T_m_ or no change was observed (Figs. 4A-F), except in the cases when carmofur binds to SARS-CoV-2 M^pro^ and EV-D68 3C^pro^, in which a T_m_ increase of 4.76 °C, and 0.87 °C was observed, respectively (Figs. 4A and 4F). A negative T_m_ shift means binding of a small molecule to a protein leads to the destabilization.

**Figure 4.**
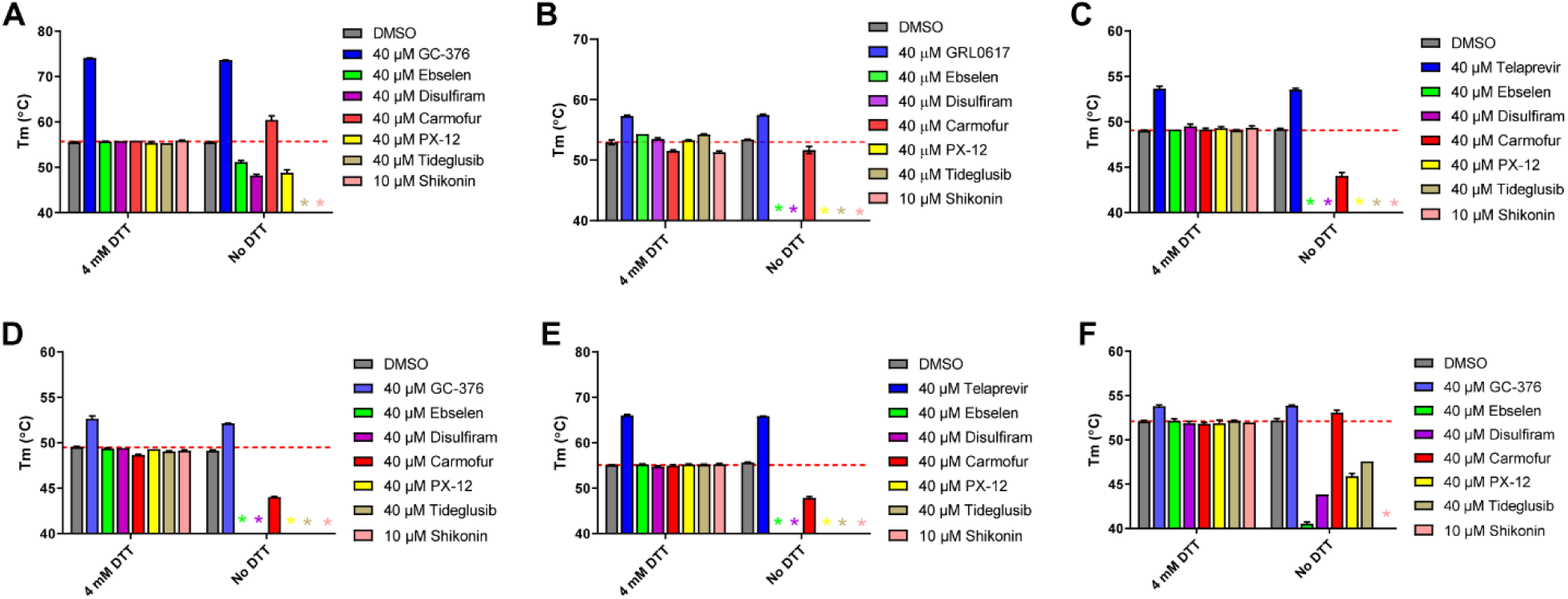
Thermal shift binding assay of SARS-CoV-2, EV-A71, and EV-D68 proteases against inhibitors investigated in this study. (**A**) SARS-CoV-2 M^pro^; (**B**) SARS-CoV-2 PL^Pro^; (**C**) EV-A71 2A^pro^; (**D**) EV-A71 3C^pro^; (**E**) EV-D68 2A^pro^; and (**F**) EV-D68 3C^pro^. 3 µM protease in its corresponding enzymatic reaction buffer in the presence of 4 mM DTT or in the absence of DTT was pre-incubated with DMSO or 40 µM protease inhibitors (shikonin was tested at 10 µM because 40 µM Shikonin completely quenches SYPRO orange dye fluorescence signal) at 30 °C for 30 min. The melting temperature (T_m_) was calculated as the mid log of the transition phase from the native to the denatured protein using a Boltzmann model.^32^ * indicates that a fluorescence peak was not observed in the melting curve; red dash line shows the protease T_m_ with DMSO in the presence of 4 mM DTT.

Taken together, when 4 mM DTT was present in the assay buffer, there was no binding between the six viral cysteine proteases and the six compounds ebselen, disulfiram, carmoufur, PX-12, tideglusib, and shikonin; without DTT, these compounds appear to non-specifically bind to these proteases, leading to destabilization.

### Ebselen, disulfiram, PX-12, tideglusib, and shikonin did not bind to SARS-CoV-2 M^pro^, while carmofur showed binding in the presence of DTT in the native mass spectrometry binding assay

To corroborate the results from the thermal shift binding assay, we next performed native mass spectrometry-based binding assays. Native MS analysis revealed that SARS-CoV-2 M^pro^ forms a dimer that was measured to have a mass of 67,595 Da (Fig. 5A). A small abundance of monomer was measured with a mass of 33,796 Da, but the intact dimer was the predominant signal (data not shown). The addition of GC376 revealed that up to two ligands bound per dimer (Fig. 5B), suggesting a binding ratio of one drug per monomer, which is consistent with its mechanism of action revealed by X-ray crystallography.^11^ The addition of 4 mM DTT shifted the equilibrium to one ligand per dimer (Fig. 5C). When carmofur was added to SARS-CoV-2 M^pro^, it bound up to three ligands per dimer, with the signal for the two bound per dimer being the most abundant (Fig. 5J). When 4 mM DTT was added it disrupted ligand binding of carmofur, with the signal for the dimer without ligand bound being the most abundant (Fig. 5K). Nevertheless, signals corresponding to one ligand per dimer and two ligands per dimer could still be detected, suggesting carmofur has moderate binding towards SARS-CoV-2 M^pro^ even in the presence of DTT (Fig. 5K). This result is consistent with our enzymatic assay result, which showed that carmofur inhibits M^pro^ with an IC_50_ of 28.2 ± 9.5 µM in the presence of 4 mM DTT (Fig. 2A). Taken together, the inhibition of M^pro^ by carmofur has certain degree of specificity, although the potency is moderate. Similarly, disulfram, ebselen, and PX-12 bound up to three, four, and five ligands per dimer respectively (Figs. 5F, 5D, and 5N) in the absence of DTT, and the addition of 4 mM DTT completely disrupted this ligand binding (Figs. 5G, 5E, and 5O), and only the dimer signal without ligand was detected. The complete disruption of this ligand binding with the addition of DTT suggests that these compounds bind non-specifically to SARS-CoV-2 M^pro^. Shikonin was found to bind up to four ligands per dimer to SARS-CoV-2 M^pro^ (Fig, 5L), and this binding was completely disrupted upon addition of 4 mM DTT (Fig. 5M). Tideglusib did not bind to M^pro^ either in the absence or the presence of 4 mM DTT at concentrations ranging from 10 µM to 40 µM (Figs. 5H-I).

**Figure 5.**
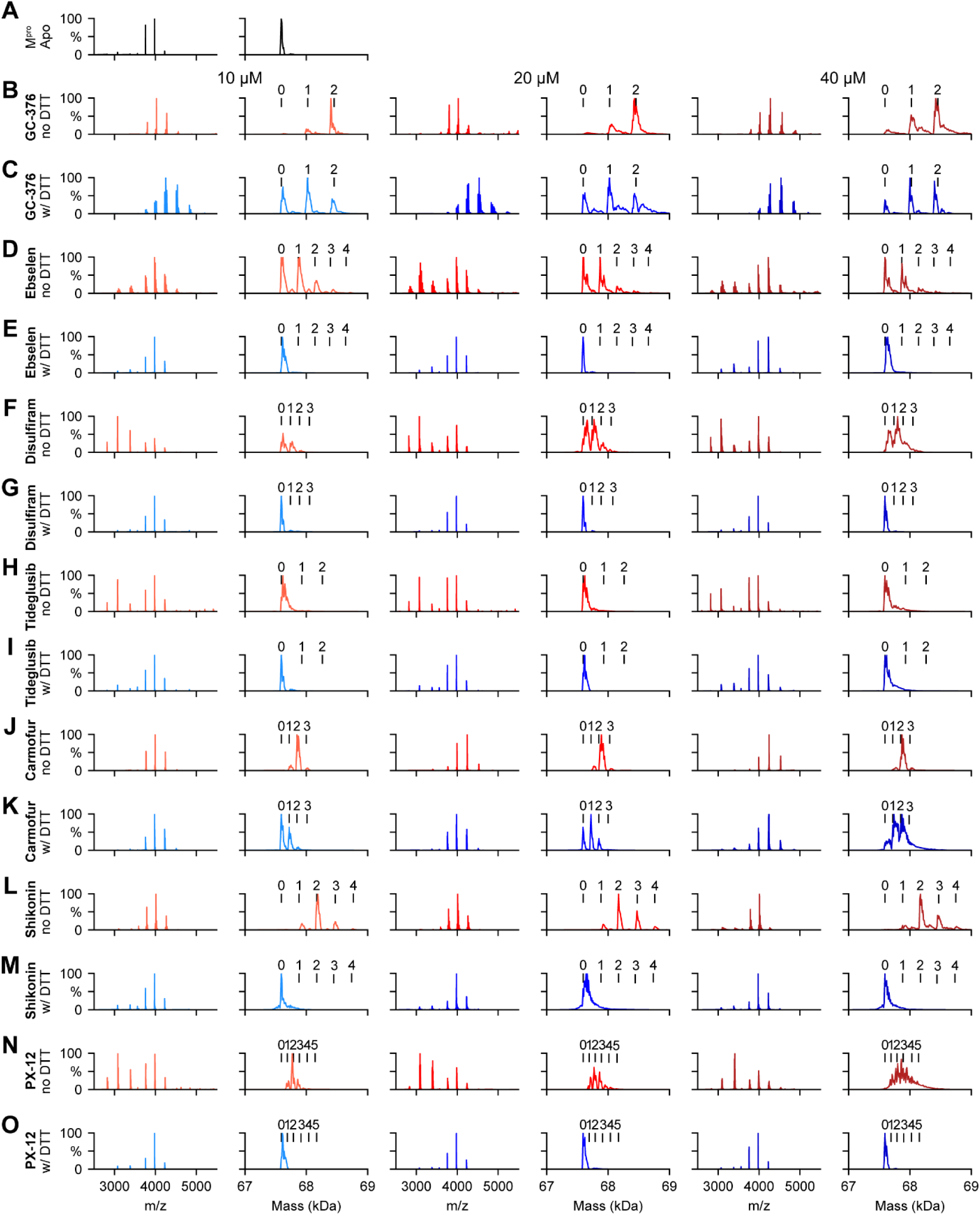
Native MS binding assay of SARS-CoV-2 M^pro^ to different protease inhibitors investigated in this study. The native mass spectra (columns 1, 3, and 5) and deconvolved mass distributions (columns 2, 4, and 6) of SARS-CoV-2 M^Pro^ without added compound (**A**) and with added GC376 (**B** and **C**), ebselen (**D** and **E**), disulfiram (**F** and **G**), tideglusib (**H** and **I**), carmofur (**J** and **K**), shikonin (**L** and **M**), and PX-12 (**N** and **O**). Spectra are shown without DTT (**B, D, F, H, J, L**, and **N**) and with 4 mM DTT (**C, E, G, I, K, M**, and **O**) and for the drug concentration of 10 µM (columns 1 and 2), 20 µM (columns 3 and 4), and 40 µM (columns 5 and 6). Dimer, one drug bound dimer, two-drug bound dimer, three-drug bound dimer, four-drug bound dimer, and five-drug bound dimer were labeled as 0, 1, 2, 3, 4, and 5 respectively.

The observation that ebselen, disulfiram, carmofur, skikonin, and PX-12 can bind to M^pro^ with more than two ligands per dimer in the absence of DTT indicates that these compounds might not only modify the catalytic cysteine C107, but also possibly bind to either allosteric sites or covalently modify other cysteine residues on M^pro^.

### Molecular dynamics simulations of the binding of M^pro^ to GC-376, carmofur, and ebselen

We performed MD simulations to compare the stability of the binding interactions identified in the X-ray structures of SARS-CoV-2 M^pro^ in complex with GC376 and carmofur. The MD simulations of M^pro^ with ebselen were carried using the highest scored docking pose.

Our previous study showed that when GC376 binds to the SARS-CoV-2 M^pro^, the covalent thioketal adduct can adapt both the S- and R-configuration.^11^ The MD simulations showed that the complexes formed with GC376 in either the S- or R-configuration did not deviate from the starting structures, which is the X-ray structure, with an RMSD in protein and ligand positions from the X-ray structure smaller than c.a. 2 Å for the protein and c.a. smaller than 2.3 Å for the ligand (Figs. 6C and 6F). The MD simulations verified stabilizing interactions observed also in the X-ray structure, which remain stable inside the binding cavity of SARS-CoV-2 as shown in the frequency interaction plot in Figs. 6A and 6D. From the protein-ligand contact plots of the GC-376 in the S-configuration (Figs. 6A-B), it is shown that it forms multiple hydrogen bonds, i.e., (a) between thiohemiketal P1 hydroxyl group and C145 peptidic NH; (b) between 2-pyrrolidone’s NH at the P1 and side chain of E166, and between peptidic NH at P1 with H164 side chain imidazole; (c) between carbamate CO, NH and benzyloxy oxygen at P2 with peptidic NH at M165, Q189 side chain CO and Q189 side chain NH, respectively. In the X-ray structure the pyrrolidone’s CO forms a hydrogen bond with H163 side chain imidazole; the MD simulation plot in Fig. 6A represents an average description of the interactions from an ensemble and not from a single snapshot in the X-ray structure. The polar 2-pyrrolidone group is oriented towards the solvent exposed S1 pocket while isobutyl group at P2 position is oriented comfortably towards the hydrophobic S2 site formed by H41, M49, and M169, while the benzyloxy group facing towards L167 moves freely in the area between Q189, A191, Q192, L167, P168 (Fig. 6B). In R-configuration GC-376 is stabilized through hydrogen bonding interaction with C145, H164, E166, Q189 and also G143 and H41, but the hydrogen bond interactions frequency is less when pyrrolidone side chain is buried in S1 pocket (Figs. 6D-E).

**Figure 6.**
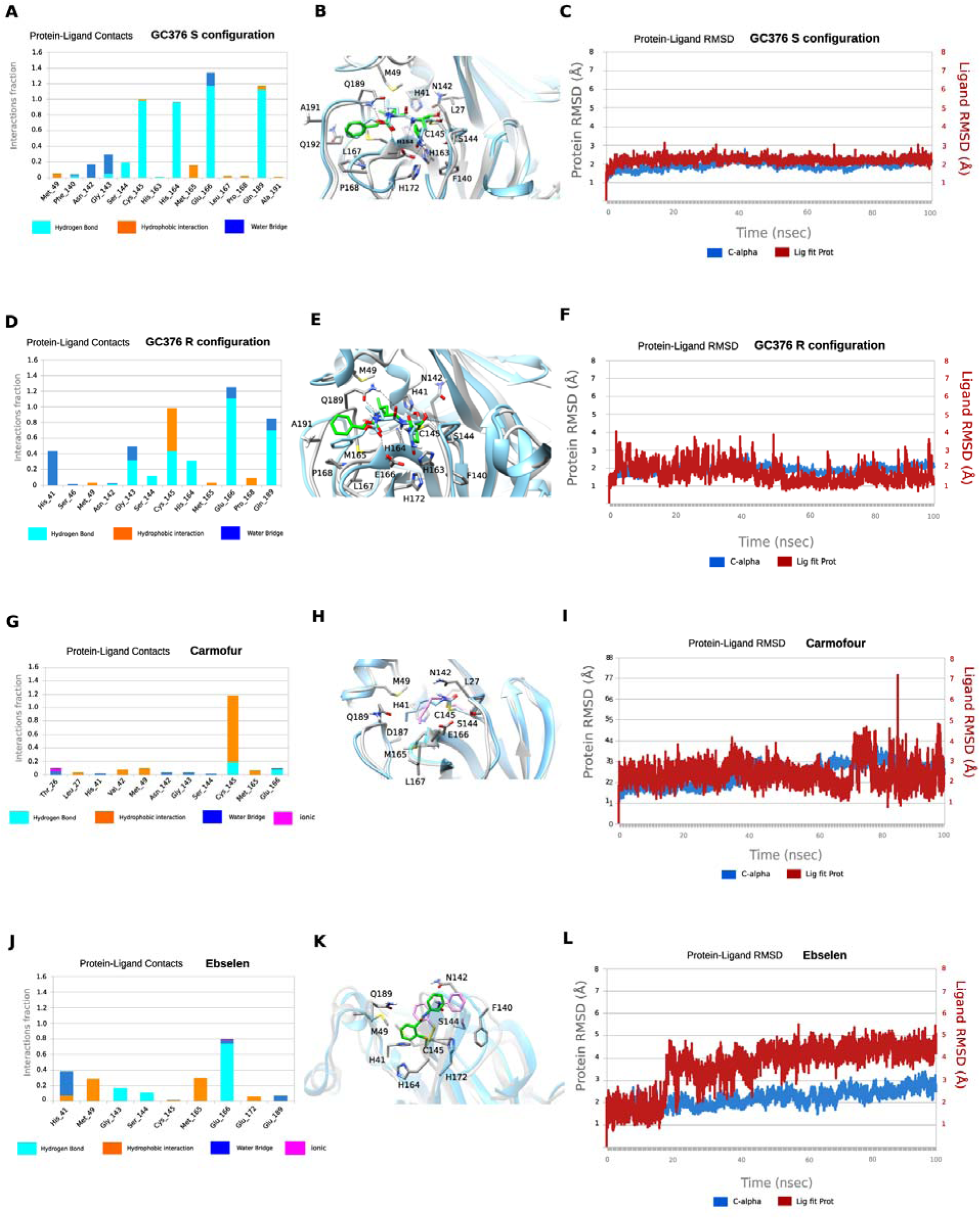
MD simulations of SARS-CoV-2 M^pro^ with its inhibitors. In **A, D, G**, and **J** hydrogen bonding interactions bar is depicted in light blue, van der Waals in orange, water bridges in blue, and ionic interactions in magenta. Interactions are plotted from 100-ns MD simulations for the complexes between the covalently bound GC376-S, GC376-R, carmofur and ebselen inside SARS-CoV-2 M^pro^. They are considered important when frequency bar is ≥ 0.2. In **B, E, H**, and **K** the last snapshots of the above mentioned 100ns-MD simulated complexes were overlaid with experimental structures with PDB IDs 6WTT for GC376-S, GC376-R and 7BUY for carmofur and a covalent docking pose for ebselen. In **C, F, I**, and **L** the RMSD plots of Cα carbons (blue diagram, left axis) and of ligand (red diagram, right axis) of the above mentioned 100ns-MD simulated complexes. The starting structures are the experimental determined structures with PDB IDs of 6WTT GC376-S, GC376-R and 7BUY for carmofur and a covalent docking poses for ebselen.

In contrast to the numerous stabilizing interactions of GC-376 in both the S- and R-configurations, carmofur interacts mainly with C145 through van der Waals and hydrogen bonding interactions (Figs. 6G-H). In carmofur the MD simulations show that the RMSD in protein and ligand positions from the X-ray structure are both c.a. 3 Å for the protein and the ligand. The hexyl side chain is oriented inside the binding cavity, from S1 to S3, but the interactions of the drugs cannot be specific for M^pro^ without directing hydrogen bond interactions and van der Waals complementary to the cavity. Ebselen interacts through hydrogen bonding interactions mainly with E166 and van der Waals interactions with M49 and M165 in the hydrophobic S2 binding area. Similarly to carmofur these interactions are not adequate to trap effectively the small drug inside the wide binding area of M^pro^ (Figs. 6J-K), and the drug can rotate around the phenyl-CO bond, resulting a high RMSD of c.a 5.4 Å (Fig. 6L).

Compared to GC376, the protein-ligand contact plots of carmofur and ebselen suggest that these two compounds bind to SARS-CoV-2 M^pro^ with reduced affinity, corroborating with their non-specific inhibition mechanism.

### Ebselen, disulfiram, carmofur, PX-12, tideglusib, and shikonin had no cellular antiviral activity against EV-A71 and EV-D68

If the enzymatic inhibition potency IC_50_ values obtained in the absence of DTT can be used to faithfully predict the cellular antiviral activity, one would expected all six compounds ebselen, disulfiram, carmofur, PX-12, tideglusib, and shikonin will have potent antiviral activity against EV-A71 and EV-D68. To test this hypothesis, the cellular antiviral activity of ebselen, disulfiram, carmofur, PX-12, tideglusib, and shikonin against EV-A71 and EV-D68 viruses were tested in RD cells using the viral cytopathic effect (CPE) assay.^33^ GC376 and telaprevir were included as positive controls as 3C^pro^ and 2A^pro^ inhibitors. GC376 inhibited EV-A71 and EV-D68 with EC_50_ values of 0.2 µM and 0.9 µM, respectively (Table 2). Telaprevir inhibited EV-D68 with an EC_50_ of 0.4 µM (Table 2). However, none of the six compounds showed antiviral activity against either EV-A71 or EV-D68 at the highest non-toxic drug concentration (Table 2).

**Table 2.** Cellular antiviral assay results of ebselen, disulfiram, carmofur, PX-12, tideglusib, and shikonin against EV-A71 and EV-D68.

|  | EV-A71<br>CPE assay<br>EC<sub>50</sub> (μM)/CC<sub>50</sub> (μM)<sup>a</sup> | EV-D68<br>CPE assay<br>EC<sub>50</sub> (μM)/CC<sub>50</sub> (μM) |
| --- | --- | --- |
| GC-376 | 0.2 ± 0.1/>50 | 0.9 ± 0.0/>50 |
| Telaprevir | N.T. | 0.4 ± 0.1/48.8 ± 4.1 |
| Ebselen | >20/17.0 ± 0.70 | >10/5.4 ± 0.2 |
| Disulfiram | >10/8.3 ± 0.6 | >3/1.5 ± 0.1 |
| Tideglusib | >20/16.6 ± 1.2 | >20/12.8 ± 0.7 |
| Carmofur | >50/47.2 ± 4.8 | >20/18.5 ± 1.5 |
| Shikonin | >1/0.8 ± 0.0 | >1/0.4 ± 0.0 |
| PX-12 | >10/7.1 ± 0.5 | >20/16.5 ± 2.4 |

<sup>a</sup>EC<sub>50</sub> and CC<sub>50</sub> (μM) = mean ± standard deviation. The values are the mean ± standard deviation from three replicates.

## Discussion

To combat COVID-19 pandemic, researchers around the globe are racing to come up with effective countermeasures. Promising progress has been made in developing vaccines and antiviral drugs. Antivirals are necessary complements of vaccines and are needs for post-infection treatment. Among the viral proteins under investigation as drug targets for SARS-CoV-2, the viral RdRp is the most extensively studied, which was followed by the viral M^pro^.^35^ Antiviral drug discovery targeting M^pro^ started with the initial efforts of developing of inhibitors against rhinovirus 3C protease (3C^pro^). The rhinovirus 3C^pro^, the enterovirus 3C^pro^, human norovirus 3CL protease, and the coronavirus 3CL protease (M^pro^) all share the same substrate preference for the glutamate at the P1 position, suggesting that 3C^pro^ or 3CL^pro^ inhibitors are promising drug candidates for broad-spectrum antivirals. Over the past few decades, significant progress has been made in designing 3C^pro^ or 3CL^pro^ inhibitors. Rupintrivir (AG7088) and AG7404 are prominent examples of human rhinovirus 3C^pro^ inhibitors that have been evaluated in clinical trials for the treatment of rhinovirus infection. For coronaviruses, GC376 is one of the most advanced lead compounds. It showed broad-spectrum in vitro antiviral activity against SARS-CoV and MERS-CoV, and in vivo antiviral activity in cats infected with feline infectious peritonitis virus.^36-37^ Given the sequence similarity between SARS-CoV-2 and SARS-CoV M^pro^, it became apparent that existing 3C^pro^ or 3CL^pro^ inhibitors might be active against SARS-CoV-2 M^pro^. Indeed, the most potent M^pro^ inhibitors reported so far such as N3, 13a, 13b, and GC376 all contain the pyrrolidone substitution in the P1 position, with variations in the reactive warhead and P2, P3, and P4 substitutions.^11-14^ Interestingly, six compounds ebselen, disulfiram, carmofur, PX-12, tideglusib, and shikonin, which share no structural similarity with GC376, were claimed as novel SARS-CoV-2 M^pro^ inhibitors.^12^ MS/MS analysis revealed that ebselen, PX-12, and carmofur were able to covalently modify the catalytic cysteine C145 of SARS-CoV-2 M^pro^.^12^

In line with these documented polypharmacology of ebselen, disulfiram, carmofur, PX-12, tideglusib, and shikonin, we are interested in validating these compounds against SARS-CoV-2 M^pro^ inhibition. Our enzymatic assay results showed that the inhibition of SARS-CoV-2 M^pro^ by these six compounds is dependent on the reducing reagent DTT. In the absence of DTT, all six compounds ebselen, disulfiram, carmofur, PX-12, tideglusib, and shikonin not only showed potent inhibition against M^pro^, but also two related viral proteases the EV-A71 and EV-D68 3C^pro^, as well as three unrelated viral proteases the SARS-CoV-2 PL^pro^, the EV-A71 and EV-D68 2A^pro^ (Fig. 2). However, upon addition of 4 mM DTT, the broad-spectrum enzymatic inhibition of these compounds was largely diminished, except carmofur and tideglusib, which had weak inhibition against M^pro^ and PL^pro^ with IC_50_ values of 28.2 and 30.4 µM, respectively (Fig. 2; Table 1). In line with the enzymatic assay results, thermal shift binding assay and native MS assay showed that ebselen, disulfiram, PX-12, tideglusib, and shikonin did not bind to M^pro^ in the presence of DTT, while carmofur could still bind to M^pro^ with the addition of DTT (Figs. 4-5). These results suggest that the inhibition of M^pro^ by carmofur has certain specificity, although the potency is relatively weak. In contrast, the inhibitory effect and binding of control compounds GC376 against M^pro^, EV-A71 and EV-D68 3C^pro^, GRL0618 against PL^pro^, and telaprevir against EV-A71 and EV-D68 2A^pro^ were not affected by the addition of DTT (Figs. 2, 4, 5). MD simulations provided additional evidence showing that the drug-bound M^pro^ complex is more stable for specific inhibitor GC376 than promiscious compounds ebselen and carmofur (Fig. 6). Furthermore, it is generally assumed that for specific inhibitors, the enzymatic inhibition potency IC_50_ value could be used to predict the cellular antiviral activity. However, despite their apparent inhibition of the EV-A71 and EV-D68 2A^pro^ and 3C^pro^ in the absence of DTT (Table 1), none of the six compounds ebselen, disulfiram, PX-12, tideglusib, and shikonin showed cellular antiviral activity against EV-A71 and EV-D68 in the CPE assay (Table 2). Therefore, caution should be taken when interpreting the enzymatic assay inhibition IC_50_ values of cysteine proteases obtained in the absence of reducing reagent such as DTT. In the absence of DTT, the apparent inhibition might be due to either alkylation or oxidation of the cysteine residue by reactive compounds. To rule out such non-specific effect, reducing reagents such as DTT, β-ME, or GSH should be added to the enzymatic buffer. Specific cysteine protease inhibitors should not show significant IC_50_ shift upon addition of reducing reagent. Moreover, counter screening against unrelated cysteine proteases should also be performed as a secondary assay to confirm the specificity.

## Materials and Methods

### Cell lines and viruses

Human rhabdomyosarcoma (RD) cells were maintained in Dulbecco’s modified Eagle’s medium (DMEM), supplemented with 10% fetal bovine serum (FBS) and 1% penicillin-streptomycin antibiotics. Cells were kept at 37°C in a 5% CO_2_ atmosphere. EV-D68 strain US/MO/14-18947 (ATCC NR-49129) was purchased from ATCC and amplified in RD cells prior to infection assays. EV-A71 strain 5865/SIN/000009 was obtained from Dr. Chan at the Department of Medical Microbiology, Faculty of Medicine, University of Malaya.^38^

### Protein expression and purification

SARS-CoV-2 M^pro^: SARS-CoV-2 M^pro^ gene from strain BetaCoV/Wuhan/WIV04/2019 in the pET29a(+) vector with E. coli codon optimization was ordered from GenScript (Piscataway, NJ). The expression and purification of SARS-CoV-2 M^pro^ was described as previously.^11^

SARS-CoV-2 PL^pro^: SARS-CoV-2 papain-like protease (PL^pro^) gene (ORF 1ab 1564 to 1876) from strain BetaCoV/Wuhan/WIV04/2019 with E. coli codon optimization was ordered from GenScript in the pET28b(+) vector. pET28b(+) plasmid with SARS-CoV-2 PL^pro^ gene was transformed into BL21(DE3) cells with kanamycin selection. A single colony was picked to inoculate 10 ml LB media and was cultured 37 °C overnight. This 10 ml culture was added to 1 liter LB media and grown to around OD600 of 0.8. This culture was cooled on ice for 15 min, then induced with 0.5 mM IPTG. Induced cultures were incubated at 18°C for an additional 24 h and then harvested, lysed same way as SARS-CoV-2 M^pro^ protein.^11^ The supernatant was incubated with Ni-NTA resin for overnight at 4 °C on a rotator. The Ni-NTA resin was thoroughly washed with 30 mM imidazole in wash buffer (50 mM Tris [pH 7.5], 150 mM NaCl, 2 mM DTT), then PL^pro^ protein was eluted from Ni-NTA with 300 mM imidazole. Eluted PL^pro^ was dialyzed against 100-fold volume dialysis buffer (50 mM Tris [pH 7.5], 150 mM NaCl, 2 mM DTT) in a 10,000-molecular-weight-cutoff dialysis tubing.

EV-A71 2A^pro^: EV-A71 2A^pro^ gene from strain EV-A71/7D3 (genbank accession number MF973167) with E. coli codon optimization was ordered from GenScript in the pET28b(+) vector. The expression and purification of EV-A71 2A^pro^ is same as SARS-CoV-2 PL^pro^ as described in the above section.

EV-A71 3C^pro^: EV-A71 3C^pro^ gene from strain EV-A71/7D3 (genbank accession number MF973167) with E. coli codon optimization was ordered from GenScript in the pET28b(+) vector. The expression and purification of EV-A71 3C^pro^ is same as SARS-CoV-2 PL^pro^ as described in the above section.

EV-D68 2A^pro^: The EV-D68 2A^pro^ gene from strain US/KY/14-18953 with E. coli codon optimization was ordered from GenScript in the pET28b(+) vector. The expression and purification of EV-D68 2A^pro^ was described as previously.^33^

EV-D68 3C^pro^: EV-AD68 3C^pro^ gene from strain US/KY/14-18953 with E. coli codon optimization was ordered from GenScript in the pET28b(+) vector. The expression and purification of EV-D68 3C^pro^ is same as SARS-CoV-2 PL^pro^ as described in the above section.

### FRET substrate Peptide synthesis

The FRET-based peptide substrates used for the enzymatic assay were shown below:

SARS-CoV-2 M^pro^ substrate: Dabcyl-KTSAVLQ/SGFRKME-Edans

SARS-CoV-2 PL^Pro^ substrate: Dabcyl-FTLRGG/APTKV-Edans

EV-A71 2A^pro^ substrate: Dabcyl-TAITTL/GKFGQE-Edan

EV-A71 3C^pro^ substrate: Dabcyl-IEALFQ/GPPKFRE-Edan

EV-D68 2A^pro^ substrate: Dabcyl-KIRIVNT/GPGFGGE-Edan

EV-D68 3C^pro^ substrate: Dabcyl-KEALFQ/GPPQFE-Edans

The synthesis of SARS-CoV-2 M^pro^, PL^pro^, EV-A71 2A^pro^, EV-D68 2A^pro^ and EV-D68 3C^pro^ substrates were described previously.^11, 33^

### Enzymatic assays

The IC_50_ values of the testing compounds against various SARS-CoV-2, EV-A71 and EV-D68 proteases in the presence or in the absence of 4 mM DTT were measured with a common protocol as the following: 100 µl protease (SARS-CoV-2 M^pro^ at 100 nM; SARS-CoV-2 PL^pro^ at 200 nM; EV-A71 2A^pro^ at 3 µM; EV-A71 3C^pro^ at 2 µM, EV-D68 2A^pro^ at 1 µM, or EV-D68 3C^pro^ at 100 nM) was incubated with various concentrations of testing inhibitors at 30°C for 30 min in its reaction buffer in 96-well plate, and then the reaction was initiated by adding FRET substrate (SARS-CoV-2 M^pro^ and PL^pro^ substrates at 10 µM; and EV-A71 and EV-D68 substrates at 20 µM), the reaction was monitored for 2□h, and the initial velocity was calculated using the data from the first 15□min by linear regression. The IC_50_ was calculated by plotting the initial velocity against various concentrations of testing inhibitor by using a four parameters dose-response curve in Prism (v8.0) software.

SARS-CoV-2 M^pro^ reaction buffer: 20 mM HEPES, pH 6.5, 120 mM NaCl, 0.4 mM EDTA, and 20% glycerol;

SARS-CoV-2 PL^pro^ reaction buffer: 50 mM HEPES, pH7.5, 0.01% triton X-100;

EV-A71 2A^pro^ reaction buffer: 50 mM Tris pH 7.0, 150 mM NaCl, 10% glyceol;

EV-A71 3C^pro^ reaction buffer: 50 mM Tris pH 7.0, 150 mM NaCl, 1 mM EDTA, 10% glycerol;

EV-D68 2A^pro^ reaction buffer: same as EV-A71 2A^pro^ reaction buffer;

EV-D68 3C^pro^ reaction buffer: same as EV-A71 3C^pro^ reaction beffer.

### The thermal shift binding assay (TSA)

The thermal shift binding assay (TSA) was carried out using a Thermal Fisher QuantStudio™ 5 Real-Time PCR System as described previously.^32-33^ Briefly, 3 µM protease in its enzymatic reaction buffer (see the above Enzymatic assays section for the reaction buffer components) in the presence of 4 mM DTT or in the absence of DTT, was incubated with testing compounds at 30 °C for 30 min in 96-well PCR plate. 1X SYPRO orange dye was added and fluorescence of the well was monitored under a temperature gradient range from 20 °C to 90 °C with 0.05 °C/s incremental step. The melting temperature (T_m_) was calculated as the mid-log of the transition phase from the native to the denatured protein using a Boltzmann model (Protein Thermal Shift Software v1.3).

### Native Mass Spectrometry

The native MS binding assay of SARS-CoV-2 M^pro^ was performed using previously described methods.^11^ Briefly, purified SARS-CoV-2 M^pro^ was buffer exchanged into 0.2 M ammonium acetate (pH 6.8) at a protein concentration of 6 μM. Each of the ligands tested (GC376, ebselen, disulfiram, tideglusib, carmofur, shikonin, and PX-12) was diluted to 200 and 100 μM in ethanol. The compounds were then titrated into the protein sample to give a final drug concentration of 10 µM, 20 µM, or 40 µM. For the ligand binding studies containing dithiothreitol (DTT), a 40 mM stock of DTT was dissolved in water. A final concentration of 4 mM DTT was added to each of those samples. For the ligand binding studies without DTT added, an equal volume of nanopore water was added to the samples in place of DTT. The final concentration of protein in each of the samples was 4.9 μM. Each sample contained 4.5 μL of protein, 0.5 μL of ligand, and 0.5 μL of DTT or water. The samples were mixed and incubated at room temperature for 30 minutes prior to analysis.

Native mass spectrometry (MS) was performed as previously described using Q-Exactive HF quadrupole-Orbitrap mass spectrometer with the Ultra-High Mass Range (UHMR) research modifications (Thermo Fisher Scientific). All of the samples were ionized in positive ion mode using 0.9 kV capillary voltage with the temperature set to 200 °C. The resolution of the instrument was set to 15,000 for all samples except for samples containing the compound Jun8-38-3, for which the resolution was set to 30,000. The trapping gas pressure within the instrument was set to 3. 50 V of source fragmentation was applied for each of the samples to aid in desolvation of the sample. All samples were analyzed between a 500-15,000 *m/z* range. All of the data were deconvolved and analyzed using UniDec.^39^

### Cytopathic Effect Assay (CPE)

The EC_50_ and CC_50_ values for the protease inhibitors investigated in this study were measured using RD cells as described previously.^32^ Briefly, RD cells were seeded and grown overnight to ∼90% confluence in 96-well plate at 37 °C 5% CO_2_ incubator. For EV-D68 virus infection, cells were washed with PBS saline and infected with virus diluted in DMEM medium with 2% FBS and 30 mM MgCl_2_. Viruses were incubated with cells for 1 hr at 33 °C followed by addition of various concentrations of testing protease inhibitors in DMEM medium with 30 mM MgCl_2_. For EV-A71 virus infection, the procedures are identical to those for EV-D68 virus, except that 30 mM MgCl_2_ was omitted in all the media and viruses were infected and incubated at 37 °C instead of 33 °C. 3 days after infection, cells were stained with 66 µg/mL of neutral red dye for 2 hr and neutral red uptake was measured at absorbance at 540 nM. The CC_50_ was measured similarly but in the absence of viral infection.

## Authors

**Chunlong Ma** – Department of Pharmacology and Toxicology, College of Pharmacy, The University of Arizona, Tucson, Arizona 85721, United States

**Yanmei Hu** – Department of Pharmacology and Toxicology, College of Pharmacy, The University of Arizona, Tucson, Arizona 85721, United States

**Julia Alma Townsend** – Department of Chemistry and Biochemistry, College of Arts and Sciences, The University of Arizona, Tucson, Arizona 85721, United States

**Panagiotis Lagarias** – Department of Pharmaceutical Chemistry, Faculty of Pharmacy, National and Kapodistrian University of Athens, Greece

**Michael Thomas Marty** – Department of Chemistry and Biochemistry, College of Arts and Sciences, The University of Arizona, Tucson, Arizona 85721, United States

**Antonios Kolocouris** – Department of Pharmaceutical Chemistry, Faculty of Pharmacy, National and Kapodistrian University of Athens, Greece

**Jun Wang** – Department of Pharmacology and Toxicology, College of Pharmacy, The University of Arizona, Tucson, Arizona 85721, United States

## Author Contributions

C. M. performed the enzymatic and thermal shift binding assays. Y. H. performed the EV-A71 and EV-D68 antiviral assays. J. A. T. performed the native mass spectrometry binding assay under the guidance of M. T. M. P. L. performed the molecular dynamics simulations under the guidance of A. K. J. W. designed and supervised this study. C. M. and J. W. wrote the manuscript with contributions from other authors.

## Acknowledgements

This research was supported by the National Institutes of Health (NIH) (Grant AI147325) and the Arizona Biomedical Research Centre Young Investigator grant (ADHS18-198859) to J. W. J.A.T. and M.T.M. were funded by the National Institute of General Medical Sciences, NIH (Grant R35 GM128624 to M.T.M.). We thank Chiesi Hellas which supported this research (SARG No 10354) and the Hellenic State Scholarships Foundation (IKY) for providing a Ph.D fellowship to P.L. (MIS 5000432, NSRF 2014-2020). This work was supported by computational time granted from the Greek Research & Technology Network (GRNET) in the National HPC facility - ARIS - under project IDs pr002021 and pr001004).

